# High-Resolution Detection of Hidden Antibiotic Resistance with the Dilution-and-Delay (DnD) Susceptibility Assay

**DOI:** 10.1101/2025.08.11.669743

**Authors:** Muqing Ma, Minsu Kim

**Affiliations:** Department of Physics, Emory University, Atlanta, GA, 30322. U.S.A; Antibiotic Resistance Center, Emory University, Atlanta, GA, 30322. U.S.A; Graduate Division of Biological and Biomedical Sciences, Emory University, Atlanta, GA, 30322. U.S.A

## Abstract

Rising rates of antibiotic treatment failure highlight the complexity of resistance mechanisms. While conventional, genetically encoded resistance is well established, recent studies have uncovered widespread non-canonical mechanisms driven by phenotypically insensitive subpopulations hiding in seemingly susceptible populations, such as heteroresistance, persistence or adaptive resistance. These variants are not only clinically significant but also act as evolutionary precursors to fully resistant populations, rendering infections increasingly difficult to control. Yet standard antibiotic susceptibility tests lack the resolution and dynamic range needed to detect this wide spectrum of resistance mechanisms and their evolutionary progression, often misguiding clinical decisions, obscuring mechanisms of failure, and underestimating the true epidemiology of antibiotic resistance. To address these limitations, we introduce the Dilution-and-Delay (DnD) assay—a practical, high-resolution approach that implements two basic principles of bacterial growth in the antibiotic susceptibility testing format. Using well-defined synthetic communities and strains, we demonstrate that the DnD assay quantifies viable cells over more than eight orders of magnitude and detects rare antibiotic-insensitive cells at frequencies as low as 1 in 100 million. It additionally reports the standardized MIC as a secondary output, thereby capturing the average antibiotic response of the majority population and heterogeneity of minority subpopulations within a single assay. We scaled this approach for high-throughput application, classifying ∼120 previously uncharacterized clinical isolates of *Klebsiella pneumoniae, Enterobacter cloacae*, *Escherichia coli*, *Pseudomonas aeruginosa* and *Acinetobacter baumannii*—across five antibiotics. This robust, quantitative, and scalable platform opens the door to next-generation antibiotic susceptibility testing, with broad utility in basic research, clinical diagnostics, and epidemiological surveillance. This improved testing will guide evidence-based clinical decisions and prevent the rapid evolution of antibiotic resistance, thereby counteracting the rising rate of treatment failure.

**Significance:** Antibiotic treatment failure due to inappropriate prescription remains a critical threat to global health. It also accelerates the evolution of resistance, making infection increasingly difficult to control. A key underlying issue is that current antibiotic susceptibility testing lacks the resolution and dynamic range needed to detect a wide range of resistance mechanisms and their evolutionary progression, leading to misinformed clinical decisions. While population analysis profiling (PAP) offers sufficient accuracy, it is too labor-intensive for routine use. Single-cell approaches are technically demanding and resource-intensive. Here, we present a susceptibility testing strategy that preserves the operational efficiency and compatibility of conventional workflows, while dramatically extending their resolution and dynamic range. Our approach is scalable, quantitative, and well-suited for high-throughput applications in both research and clinical settings—closing a critical diagnostic gap in the fight against antibiotic treatment failure.

## Introduction

Antibiotics are foundational to modern medicine—significantly improving quality of life, extending human lifespans, and enabling complex medical procedures. However, treatment failure due to inappropriate antibiotic use remains common, worsening patient outcomes and increasing the burden on healthcare systems ^1,2^. Antibiotic susceptibility testing plays a critical role in combating this threat by guiding clinical decisions and ensuring treatments are tailored to the infecting pathogen.

Concerns about antibiotic failure date back to the very beginning of the antibiotic era, when Alexander Fleming warned that improper use of penicillin could drive the emergence of resistance—a prediction that proved prescient. Conventional antibiotic resistance arises by the acquisition of genetically encoded determinants through mutations or horizontal gene transfer ^3^, which confer population-scale advantages under antibiotic pressure.

However, recent studies have shed new light on diverse mechanisms by which bacteria evade antibiotic treatments. In hetero-resistance, the majority of cells in a genetically identical population remain susceptible, while a minority exhibits high-level resistance ^4,5^. In persistence, a subpopulation transiently enters a dormant or slow-growing state, surviving lethal antibiotic exposure and resuming growth once treatment ends ^6,7^. Sublethal antibiotic exposure can also induce adaptive resistance, allowing certain cells to gradually withstand higher drug concentrations ^8,9^. These traits can propagate through lineages and be stably maintained in the population ^10,11^. Mounting evidence indicates that these non-canonical forms of antibiotic insensitivity are not exceptions but widespread contributors to treatment failure ^12,13^.

Importantly, these antibiotic-insensitive phenotypes are often part of a broader evolutionary landscape. Heteroresistance or persistence often emerges early during the evolution under antibiotic treatments ^14-16^. Once antibiotic-insensitive subpopulations are present, inappropriate treatments can accelerate their acquisition of genetic resistance—providing an alternate route to rapid, population-wide resistance ^14-17^.

However, current antibiotic susceptibility testing methods are not adequate for detecting this full range of resistance mechanisms and their evolutionary progression. The gold-standard broth-based assay evaluates visual turbidity in liquid media containing antibiotics to find the minimum inhibitory concentration (MIC) ^18,19^—a single numerical value that summarizes the response of the entire population ^20^. Alternative methods such as disk diffusion and E-tests apply the same principle on agar plates, using the diameter of the inhibition zone to infer the MIC ^21^. In all cases, the MIC obtained is compared with the pre-defined clinical breakpoint to classify the population as susceptible or resistant ^22^.

While simple and widely adopted, these current tests average over the population, masking underlying heterogeneity. As a result, they fail to detect non-canonical forms of antibiotic insensitivity embedded within an otherwise susceptible population ^23-25^, misclassifying the population as fully susceptible. This, in turn, results in inappropriate therapy and “unexplained” treatment failures. More concerningly, when inappropriate therapy drives the evolutionary progression of minority variants to full-scale resistance ^14-17^, current tests remain blind to this process—detecting it only after resistant clones have swept through and become fixed in the population, by which point it is too late to adjust therapy and reverse the course. This gap underscores the urgent need for high-resolution susceptibility testing capable of detecting antibiotic-insensitive cells even when they occur at very low frequencies within susceptible populations.

The ideal solution would be single-cell-level detection of bacterial survival. Leveraging advances in microscopy and microfluidics, we and others have measured antibiotic susceptibility at single-cell resolution, detecting survival frequencies as low as 1 in 10,000 (10⁻⁴) ^26,27^. While promising, this range is not sufficient to reliably detect non-canonical mechanisms of antibiotic insensitivity such as heteroresistance or persistence, which can occur at frequencies far below this range (e.g., 10^-7^ for heteroresistance) ^23-25,28,29^. Moreover, these approaches depend on specialized instrumentation and technical expertise, making them inaccessible to most laboratories.

Historically, agar-based population analysis profiling (PAP) has been the method of choice for detecting heteroresistance ^23-25^. In PAP, bacterial cultures are plated across a range of antibiotic concentrations, and colonies that grow at high concentrations are counted to quantify resistant cells. Although highly sensitive, PAP is labor-intensive and time-consuming—requiring agar preparation and manual CFU counting—making it impractical for routine diagnostics.

This diagnostic blind spot obscures the diverse mechanisms underlying treatment failure, misguides clinical decision, and masks the true magnitude of antibiotic resistance, ultimately delaying the development of evidence-based clinical guidelines. There is a pressing need for susceptibility testing platforms that retain the operational efficiency of current antibiotic susceptibility testing, while offering dramatically improved resolution and dynamic range.

To address this challenge, we investigated two fundamental principles of bacterial growth, dilution-to-extinction and delay-to-growth, in the standard testing format, developing a practical, scalable, high-resolution strategy for antibiotic susceptibility testing.

## Result

The gold standard for antibiotic susceptibility testing is broth-based culture, standardized by international organizations, including the Clinical and Laboratory Standards Institute (CLSI) ^18^ or the European Committee on Antimicrobial Susceptibility Testing (EUCAST) ^19^. These protocols specify inoculating a fixed number of bacterial cells (*N*, typically ∼5×10^5^) into antibiotic-containing media and visually observing the turbidity after a defined period (*T,* typically ∼18 hours) to determine the MIC of the sample.

However, *N* and *T* are procedural parameters—not intrinsic biological properties. We hypothesized that systematically varying these parameters could reveal resistant cells hidden in a susceptible population.

### Dilution-to-extinction quantifies rare resistant cells through serial dilution

Consider a bacterial culture in which resistant cells occur at a frequency of 10⁻⁷. If the standard inoculum *N* ≍ 5×10^5^ is used, it will likely contain no resistant cells and will be sterilized by antibiotic treatment. In contrast, a culture of *N* ≈ 10⁹ cells would almost certainly include resistant cells, allowing the culture to grow to become turbid in the presence of antibiotics. Therefore, by systematically varying *N*, we can infer the abundance of resistant subpopulations based on turbidity.

We formalized this concept into a dilution-to-extinction assay (Fig. 1a). Starting from a high-density culture, we performed serial dilutions and incubated them in antibiotic-containing media. At low dilutions, resistant cells are likely present and grow, driving turbidity. As the dilution increases, the probability of including resistant cells decreases, and eventually, a dilution contains none—resulting in no growth. The first dilution in the series that lacks turbidity defines the extinction point (Fig. 1a). For example, if the original population contains tens of resistant cells, the first 10-fold dilution would still turn turbid, while the second would not. In our analysis, we determined the number of dilutions required to reach the extinction point to estimate the number of resistant cells in the original sample.

**Figure 1:**
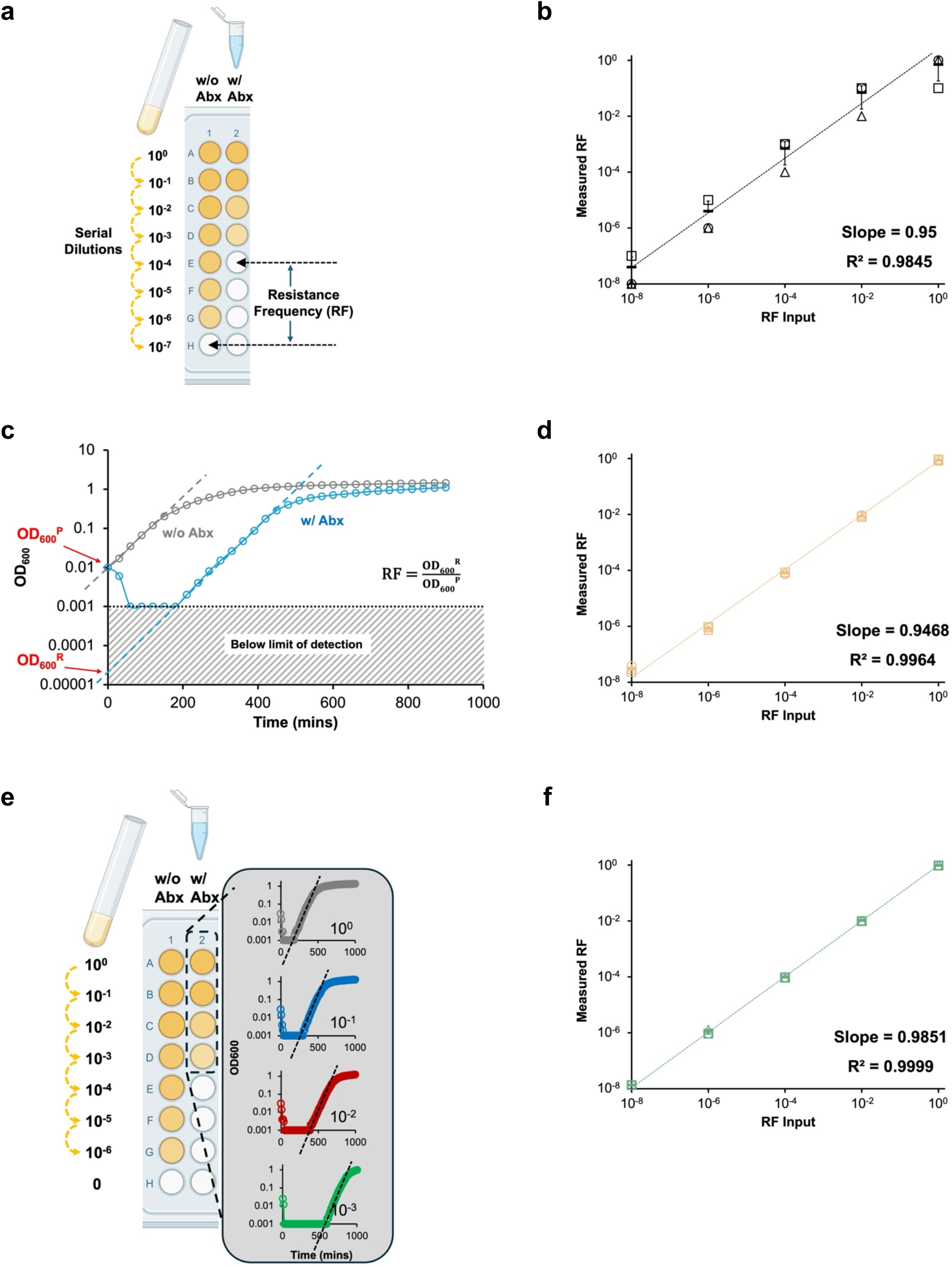
High-resolution quantification of rare antibiotic-resistant subpopulations. **a.** Dilution-to-extinction assay. A bacterial culture (10⁹ cells/mL) is serially diluted and incubated with and without antibiotics (abx) for ∼18 hours. The first dilution lacking visible turbidity defines the extinction point, which estimates the number of resistant cells in the original population. **b.** To validate this approach, a colistin-susceptible strain (AMK105) was spiked with a fully resistant strain (AMK104) at defined input frequencies. Resistance frequencies (RFs) were inferred based on extinction points (Fig. 1a). They closely matched the known inputs across several orders of magnitude. Linear regression yielded a slope near 1 and R² = 0.985. **c.** Delay-to-growth assay. In antibiotic-treated cultures, the initial OD₆₀₀ decline reflects killing of the susceptible population, followed by delayed regrowth from rare resistant cells. Exponential regrowth is extrapolated back to time zero to estimate OD₆₀₀ᴿ, representing the density of resistant cells in the original culture. Comparing this to the initial OD₆₀₀ᴾ of the total population yields an estimate of RF. **d.** RF values inferred from delay-to-growth closely matched the known input frequencies, with strong linear correlation (slope ≈ 1, R² = 0.996). **e.** Dilution-and-Delay (DnD) assay. The two methods were integrated by performing serial dilutions in a 96-well plate and continuously monitoring OD₆₀₀ in each well. Wells that remained clear defined the extinction point, while wells showing delayed regrowth were used to infer RF, as described in Fig. caption 1a and 1c. Multiple wells exhibiting delayed growth were used to calculate an average RF. The average RF values were cross-validated against extinction points to ensure internal consistency. **f.** Inferred average RF from the DnD assay was plotted against known input frequencies. The DnD method demonstrated strong quantitative agreement (slope ≈ 1) and exhibited the highest precision (R² = 0.999) among the three approaches. Each experiment was performed in biological triplicate. Individual replicates are shown as different symbols, with lines and error bars indicating the mean and standard deviation.

To validate this approach, we spiked a susceptible strain with known fractions of a fully resistant strain, preparing defined mixtures. We applied the dilution-to-extinction protocol in the presence of antibiotics to estimate resistant cells, and in parallel, performed the same dilution series without antibiotics to estimate the total number of viable cells (Fig. 1a). Comparing the extinction points across the two conditions enabled us to compute the resistance frequency (RF) in each mixture (Fig. 1a).

Across samples with resistant fractions spanning several orders of magnitude, the measured RF values closely recapitulated the known input frequencies (Fig. 1b). Replicate measurements, indicated by three different symbols, exhibited ∼10-fold variation, consistent with the resolution limit imposed by 10-fold serial dilution. Nonetheless, the data showed a strong linear relationship, with a slope near 1 and an R² of 0.98 (Fig. 1b). Notably, the method reliably detected resistant cells at frequencies as low as ∼10⁻⁸—well below a conventional heteroresistance threshold of ∼10⁻⁷ ^25^. This test indicates that the dilution-to-extinction assay enables unbiased detection of resistant subpopulations across a broad dynamic range.

### Delay-to-Growth Captures Rare Survivors Through Density Monitoring

The other fixed parameter in standard antibiotic susceptibility testing is the observation time (*T*), typically ∼18 hours ^18,19^. This constraint assumes that antibiotic-sensitive and -resistant populations will fully diverge within that period. However, when resistant cells are rare, their outgrowth is delayed. The less resistant cells are present, the longer this delay. The core idea behind the delay-to-growth assay is to analyze this delay under antibiotic treatment to infer the abundance of resistant cells.

We tested this principle using the defined mixtures of susceptible and resistant strains. Optical density (OD₆₀₀) was measured over time using a spectrophotometer to monitor culture growth. In the absence of antibiotics, the culture exhibited immediate exponential growth without lag (Fig. 1c, grey). In contrast, when antibiotics are present, the OD₆₀₀ initially declined due to killing of susceptible cells—often dropping below the detection limit of a spectrophotometer (Fig. 1c, cyan). This was followed by delayed increase in OD_600_, corresponding to the outgrowth of surviving resistant cells that become detectable.

To estimate the number of resistant cells in the original culture, we fit the exponential phase of the recovery curve, extrapolating it back to time zero (dashed line, Fig. 1c). This yielded an OD₆₀₀ of resistant subpopulations at *t* = 0, denoted OD₆₀₀ᴿ (Fig. 1c). To calculate its frequency relative to the total population, we compared it with the total OD_600_ of the culture (OD₆₀₀ᴾ in Fig. 1c). The resulting ratio, OD₆₀₀ᴿ / OD₆₀₀ᴾ, provides an estimate of RF in the original sample.

To assess the quantitative accuracy and dynamic range of the method, we varied the abundance of resistant cells in the defined mixture over several orders of magnitude and measured the RFs. The measured RF values closely matched the known input ratio, yielding a linear regression with a slope near 1 with R^2^= 0.996 (Fig. 1d). Therefore, the delay-to-growth approach accurately captures resistant subpopulations.

### Integrating Two Axes of Resolution: The Dilution-and-Delay (DnD) Framework

To improve both quantitative accuracy and operational efficiency, we integrated the dilution-to-extinction and delay-to-growth assays into a unified framework. Since both assays are performed in broth and rely on turbidity as the readout—whether qualitative or quantitative—they are naturally compatible. Moreover, turbidity can be monitored automatically via OD₆₀₀ measurements using standard plate readers, enabling seamless integration into high-throughput workflows. Importantly, the two assays are based on distinct underlying principles, yielding orthogonal estimates of RF. Cross-validating these independent measurements provides an internal consistency check, thereby increasing the robustness and reliability of RF quantification.

We implemented this combined strategy as the Dilution-and-Delay (DnD) assay. Following the dilution-to-extinction protocol, we prepared serial dilutions in a 96-well microtiter plate, then continuously monitored OD₆₀₀ in each well over time using a plate reader (Fig. 1e). Wells that failed to show an OD₆₀₀ increase were used to define the extinction point, while those that exhibited growth were fit to estimate RF values based on the delay-to-growth method (Fig. 1e). The dilution series also served as technical replicates, allowing us to compute an average RF across all growing wells, thereby improving statistical precision.

We validated the DnD assay using defined mixtures of fully resistant and susceptible strains. Across all input ratios, the measured RFs were within a 10-fold range of estimates based on the extinction point, providing a pleasing check of consistency (Supplementary Fig. 1). Linear regression of input versus measured RFs showed a slope near 1 and R² of 0.999 (Fig. 1f), confirming strong quantitative agreement.

### Validation Against Heteroresistant Clinical Isolates

We next applied our assays to a panel of five clinical isolates previously characterized as heteroresistant against various antibiotics ^17,30,31^. Unlike the defined artificial mixtures used above, these strains endogenously harbor resistant subpopulations with variable frequencies and resistance levels. Each strain had been previously evaluated using population analysis profiling (PAP)—a well-established method for identifying heteroresistance—revealing a gradual decline in CFUs with increasing antibiotic concentrations (red symbols in Fig. 2a-c for three strains; Supplementary Fig. 2 for the remaining two strains). This gradual decrease reflects a heterogeneous distribution of resistance within each population. By comparing CFUs at each antibiotic concentration to those in antibiotic-free conditions, we obtained RF_PAP_ (red in Fig. 2). Three replicate measurements were averaged to calculate the mean RF_PAP_: ⟨RF⟩_PAP_.

**Figure 2:**
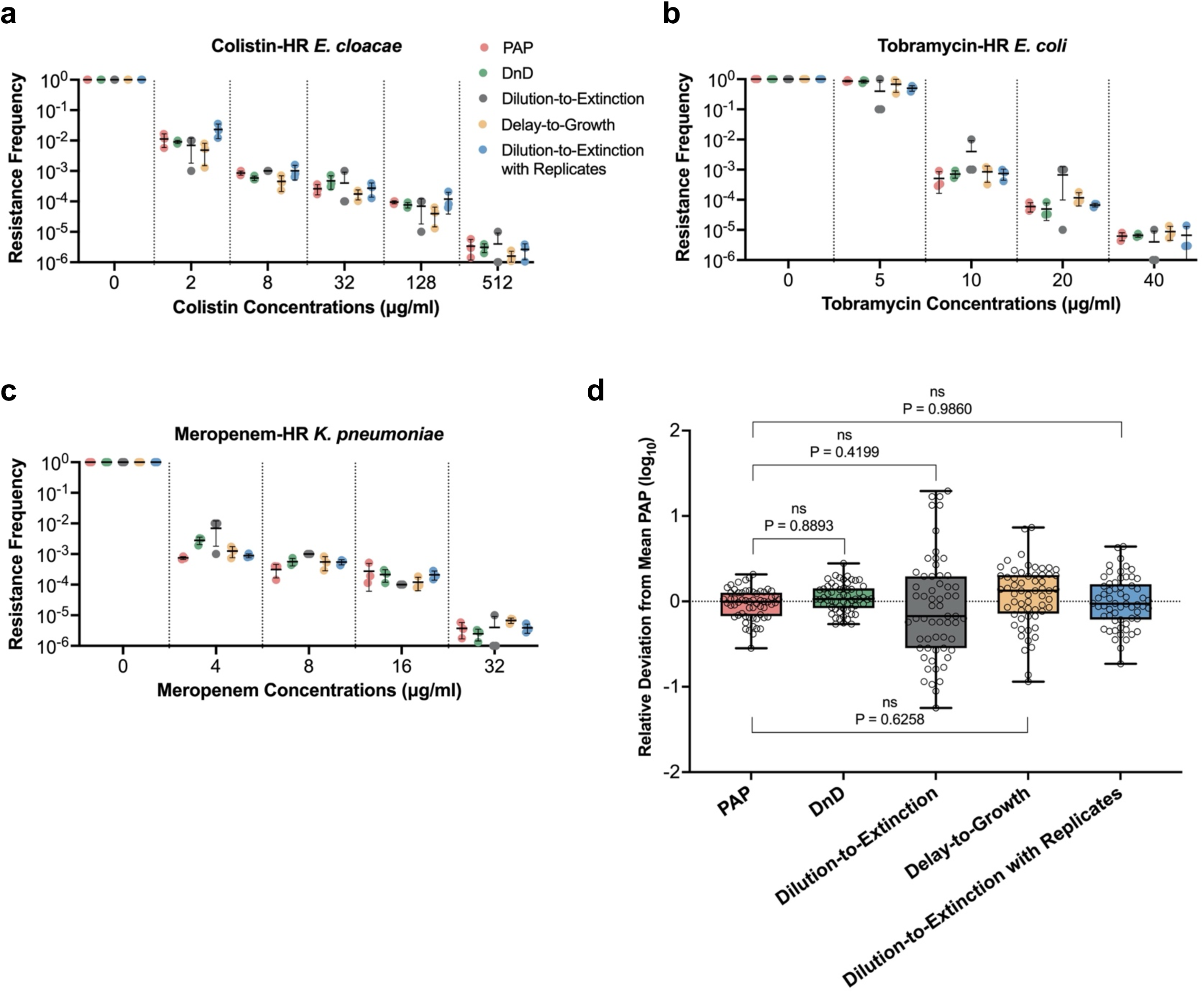
Validation of the DnD assay against heteroresistant clinical isolates. **a–c.** Resistance frequencies (RFs) were measured for five clinical heteroresistant (HR) isolates: colistin-HR *E. cloacae* (AMK107), tobramycin-HR *E. coli* (AMK117), and meropenem-HR *K. pneumoniae* (KMK4). See Supplementary Fig. 2 for two additional HR strains (AMK118, AMK120). In prior studies, population analysis profiling (PAP) was used to determine the fraction of resistant subpopulations. For direct comparison, we applied the DnD assay alongside dilution-to-extinction and delay-to-growth assays. Each assay was performed in biological triplicate; individual replicates are shown as different symbols, with black lines and error bars indicating the mean and standard deviation. All three assays recapitulated PAP-derived RF, RF_pap_, across antibiotics and concentrations. **d.** RF values from all assays (DnD, dilution-to-extinction, delay-to-growth) were statistically compared against RF_pap_. Data from panels a–c and Supplementary Fig. 2 were pooled, and each RF value was normalized to the corresponding mean PAP value (<RF>_pap_) to calculate relative deviation on a log₁₀ scale. The distribution of log₁₀(RF/(<RF>_pap_) is centered around 0, indicating strong agreement across methods. One-way ANOVA showed no statistically significant differences between methods (p > 0.05), denoted as “ns.” Notably, the DnD assay exhibited the lowest variance and a high p-value, underscoring its superior accuracy and reproducibility.

Although PAP is highly accurate, its labor- and time-intensive nature limits its utility in clinical and high-throughput settings. We therefore evaluated our DnD (green), dilution-to-extinction (grey), and delay-to-growth (yellow color) assays on the same strains. Measurements were repeated three times for each condition. All three assays produced RFs that closely matched one another and recapitulated the gradual decline observed in PAP (Fig. 2a-c and Supplementary Fig. 2).

For quantitative comparison, we characterized how our assay results deviate from that of the PAP assay by normalizing their RFs by ⟨RF⟩_PAP_, thereby quantifying their deviation by RF / ⟨RF⟩_PAP_ for each strain and antibiotic concentration. Aggregating these values (open symbols, Fig. 2d) revealed that RF values from each of our assays were centered around the ⟨RF⟩_PAP_, indicating their strong agreement. Dilution-to-extinction exhibited the most variance, about 10-fold, as discussed above. The DnD assay exhibited the lowest variance—comparable to that of PAP replicates—indicating that DnD achieves a level of precision equivalent to PAP.

We further performed statistical comparison using a one-way ANOVA test between RF values from each assay and those from PAP. All tests yielded *p* > 0.05, indicating that our assay results and PAP results are not statistically different (Fig. 2d). This finding aligns with earlier comparisons using defined mixtures, where each assay accurately captured the input frequency (Fig. 1). Notably, among the three, the DnD assay produced the highest *p*-value (approaching 1), suggesting its measurements are quantitatively indistinguishable from PAP. Together, these tests demonstrate that all three assays faithfully quantify antibiotic-resistant subpopulations, with the DnD assay providing the greatest accuracy and reproducibility.

The MIC is a widely used metric in clinical microbiology for assessing the population-scale response to antibiotics ^20^. Although the primary purpose of the DnD assay is to quantify rare antibiotic-insensitive subpopulations, it also recovers MIC values as a secondary output. By design, the DnD assay includes a dilution series that spans a wide range of inoculum sizes, including wells that approximate the standard inoculum size of ∼5 ×10^5^ cells ^18,19^. Following standard guidelines, we defined MIC as the lowest antibiotic concentration that prevented visible growth in these wells (see Supplementary Table 1 for the MIC values).

This comparison between MIC and resistance frequency (RF) underscores the fundamental detection limit of standard assays. We found that the MICs consistently corresponded to the first antibiotic concentration at which RF dropped below 10^-5^ (Fig. 2 and Supplementary Fig. 2). This detection threshold reflects the standard inoculum size in the range of 10⁵ cells. At this inoculum size, a culture with RF below 10^-5^ will no longer contain resistant cells and is cleared by antibiotics—thus reporting the MIC. As a result, conventional MIC testing can detect resistance only when resistant subpopulations are sufficiently large. However, it does not reveal the size of these subpopulations, and more critically, it fails to detect resistance when their frequency falls below the threshold.

### High-Throughput Screening of Previously Uncharacterized Clinical Isolates

In addition to quantitative accuracy, another key advantage of the DnD assay is its compatibility with automated, high-throughput screening. Because the assay is implemented in microtiter plates and utilizes optical density as a readout, it can be fully automated using a standard plate reader.

We applied the DnD assay for high-resolution detection of antibiotic resistance in a panel of ∼120 previously uncharacterized clinical isolates of *Klebsiella pneumoniae*, *Acinetobacter baumannii*, *Pseudomonas aeruginosa*, *Enterobacter cloacae*, and *Escherichia coli*. These strains were obtained from the Multi-site Gram-negative Surveillance Initiative (MuGSI), CDC. Each isolate was tested against five antibiotics: meropenem and piperacillin/tazobactam (β-lactams), tobramycin (aminoglycosides), ciprofloxacin (fluoroquinolones), and colistin (polymyxins).

For each strain and drug, we calculated the RF at the clinical breakpoint concentration; see Source Data for the raw RF values. Following a recently proposed definition for clinical screening ^14^, we classified strains into three categories: resistant (RF > 0.5), heteroresistant (10⁻⁷ < RF < 0.5), or susceptible (RF < 10⁻⁷). To validate these classifications, we independently performed PAP assays on the same isolates and antibiotics.

Out of 540 classifications, DnD and PAP matched in all but four cases, corresponding to a 99.3 % agreement rate (Table 1). The few discrepancies (highlighted in red) resulted from minor differences in RF values near classification thresholds, typically between resistant and heteroresistant categories. For instance, the strain KMK70 treated with colistin showed an RF of 58.2 % by DnD and 43.9% by PAP—both indicating high resistance, though falling on opposite sides of the 50% threshold. These mismatches reflect the use of artificial cutoff values, which can vary between studies ^23,25,30,31^, rather than substantive discrepancy between DnD and PAP tests.

**Table 1.** High-throughput Screening of ∼120 clinical isolates. *K.p*: *Klebsiella pneumoniae*, *E. coli*: *Escherichia coli*, *E. c*: *Enterobacter cloacae, P. a*: *Pseudomonas aeruginosa, A. b*: *Acinetobacter baumannii*. Following the definition in the literature for screening ^14^, we classified strains into three categories: resistant (RF > 0.5), heteroresistant (10⁻⁷ < RF < 0.5), or susceptible (RF < 10⁻⁷). See Source Data for raw RF values.

| KMK# | Species | Meropenem |  | Pip/Tazo |  | Tobramycin |  | Colistin |  | Ciprofloxacin |  | KMK# | Species | Meropenem |  | Pip/Tazo |  | Tobramycin |  | Colistin |  | Ciprofloxacin |  |
| --- | --- | --- | --- | --- | --- | --- | --- | --- | --- | --- | --- | --- | --- | --- | --- | --- | --- | --- | --- | --- | --- | --- | --- |
|  |  | PAP | DnD | PAP | DnD | PAP | DnD | PAP | DnD | PAP | DnD |  |  | PAP | DnD | PAP | DnD | PAP | DnD | PAP | DnD | PAP | DnD |
| 25 | *E. coli* | S | S | HR | HR | S | S | S | S | R | R | 48 | *K. p.* | HR | HR | HR | HR | S | S | S | S | R | R |
| 28 | *E. coli* | HR | HR | HR | HR | HR | HR | S | S | R | R | 49 | *K. p.* | R | R | R | R | S | S | S | S | R | R |
| 29 | *E. coli* | HR | HR | R | R | R | R | S | S | R | R | 50 | *K. p.* | R | R | R | R | S | S | S | S | R | R |
| 35 | *E. coli* | HR | HR | HR | HR | R | R | S | S | R | R | 51 | *K. p.* | HR | HR | R | R | S | S | S | S | R | R |
| 44 | *E. coli* | HR | HR | HR | HR | HR | HR | S | S | HR | HR | 52 | *K. p.* | HR | HR | R | R | S | S | S | S | R | R |
| 60 | *E. coli* | HR | HR | HR | HR | R | R | S | S | R | R | 54 | *K. p.* | R | R | R | R | S | S | S | S | R | R |
| 85 | *E. coli* | HR | HR | HR | HR | R | R | S | S | R | R | 55 | *K. p.* | R | R | HR | HR | S | S | S | S | R | R |
| 88 | *E. coli* | HR | HR | R | R | S | S | S | S | R | R | 57 | *K. p.* | HR | HR | R | R | HR | HR | S | S | R | R |
| 97 | *E. coli* | HR | HR | HR | HR | S | S | S | S | S | S | 58 | *K. p.* | R | R | R | R | HR | HR | S | S | R | R |
| 99 | *E. coli* | S | S | HR | HR | S | S | S | S | R | R | 59 | *K. p.* | HR | HR | R | R | R | R | S | S | R | R |
| 7 | *E. c.* | HR | HR | S | S | S | S | S | S | R | R | 61 | *K. p.* | HR | HR | R | R | R | R | S | S | R | R |
| 22 | *E. c.* | HR | HR | R | R | S | S | S | S | R | R | 62 | *K. p.* | R | R | R | R | R | R | S | S | R | R |
| 53 | *E. c.* | R | R | R | R | R | R | HR | HR | R | R | 63 | *K. p.* | R | R | R | R | S | S | S | S | R | R |
| 56 | *E. c.* | S | S | S | S | S | S | HR | HR | HR | HR | 64 | *K. p.* | R | R | R | R | S | S | S | S | R | R |
| 75 | *E. c.* | S | S | S | S | S | S | HR | HR | R | R | 65 | *K. p.* | R | R | R | R | S | S | S | S | R | R |
| 81 | *E. c.* | HR | HR | HR | HR | R | R | S | S | R | R | 66 | *K. p.* | HR | HR | R | R | S | S | S | S | S | S |
| 82 | *E. c.* | HR | HR | HR | HR | R | R | S | S | R | R | 67 | *K. p.* | R | R | R | R | S | S | S | S | R | R |
| 4 | *K. p.* | HR | HR | R | R | R | R | S | S | R | R | 68 | *K. p.* | R | R | HR | HR | R | R | S | S | R | R |
| 5 | *K. p.* | R | R | R | R | S | S | S | S | R | R | 69 | *K. p.* | R | R | HR | HR | R | R | S | S | R | R |
| 6 | *K. p.* | R | R | R | R | S | S | S | S | R | R | 70 | *K. p.* | HR | HR | HR | HR | R | R | HR | R | R | R |
| 8 | *K. p.* | HR | HR | R | R | S | S | S | S | R | R | 71 | *K. p.* | R | R | R | R | R | R | S | S | R | R |
| 9 | *K. p.* | HR | HR | R | R | HR | HR | S | S | R | R | 72 | *K. p.* | R | R | R | R | R | R | S | S | R | R |
| 10 | *K. p.* | HR | HR | HR | HR | S | S | S | S | R | R | 73 | *K. p.* | R | R | R | R | R | R | HR | HR | R | R |
| 11 | *K. p.* | HR | HR | R | R | S | S | S | S | R | R | 74 | *K. p.* | R | R | R | R | R | R | S | S | R | R |
| 12 | *K. p.* | HR | HR | R | R | S | S | S | S | R | R | 76 | *K. p.* | S | S | S | S | R | R | S | S | R | R |
| 13 | *K. p.* | HR | HR | R | R | HR | HR | S | S | HR | HR | 77 | *K. p.* | HR | HR | R | R | R | R | S | S | R | R |
| 14 | *K. p.* | R | R | R | R | S | S | S | S | HR | HR | 78 | *K. p.* | R | R | HR | HR | R | R | HR | R | R | R |
| 15 | *K. p.* | HR | HR | R | R | HR | HR | S | S | R | R | 79 | *K. p.* | HR | HR | HR | HR | R | R | S | S | R | R |
| 16 | *K. p.* | HR | HR | R | R | HR | HR | S | S | R | R | 80 | *K. p.* | HR | HR | HR | HR | R | R | S | S | R | R |
| 17 | *K. p.* | HR | HR | R | R | S | S | S | S | R | R | 83 | *K. p.* | S | S | S | S | R | R | HR | HR | R | R |
| 18 | *K. p.* | R | R | R | R | S | S | S | S | R | R | 84 | *K. p.* | HR | HR | HR | HR | R | R | S | S | R | R |
| 19 | *K. p.* | R | R | R | R | S | S | S | S | R | R | 86 | *K. p.* | HR | HR | R | R | R | R | S | S | R | R |
| 20 | *K. p.* | R | R | R | R | HR | HR | S | S | R | R | 87 | *K. p.* | R | R | R | R | R | R | S | S | R | R |
| 21 | *K. p.* | HR | HR | R | R | S | S | S | S | S | S | 89 | *K. p.* | R | R | HR | HR | S | S | HR | HR | R | R |
| 23 | *K. p.* | R | R | R | R | R | R | HR | HR | R | R | 90 | *K. p.* | R | R | HR | HR | S | S | S | S | R | R |
| 24 | *K. p.* | R | R | R | R | S | S | S | S | R | R | 91 | *K. p.* | R | R | R | R | R | R | S | S | R | R |
| 26 | *K. p.* | HR | HR | HR | HR | S | S | S | S | R | R | 92 | *K. p.* | HR | HR | R | R | R | R | S | S | R | R |
| 27 | *K. p.* | R | R | R | R | S | S | S | S | R | R | 93 | *K. p.* | R | R | R | R | R | R | S | S | R | R |
| 30 | *K. p.* | R | R | R | R | S | S | HR | HR | R | R | 94 | *K. p.* | HR | HR | HR | HR | R | R | S | S | R | R |
| 31 | *K. p.* | HR | R | HR | HR | R | R | HR | HR | R | R | 95 | *K. p.* | R | R | R | R | R | R | HR | HR | R | R |
| 32 | *K. p.* | R | R | R | R | HR | HR | S | S | R | R | 96 | *K. p.* | R | R | R | R | R | R | S | S | R | R |
| 33 | *K. p.* | HR | HR | HR | HR | R | R | HR | HR | R | R | 98 | *K. p.* | R | R | R | R | R | R | S | S | R | R |
| 34 | *K. p.* | R | R | R | R | S | S | HR | HR | R | R | 99 | *P. a.* | R | R | R | R | S | S | S | S | S | S |
| 36 | *K. p.* | R | R | R | R | S | S | S | S | R | R | 100 | *P. a.* | HR | HR | HR | HR | S | S | HR | HR | S | S |
| 37 | *K. p.* | HR | HR | R | R | R | R | HR | HR | R | R | 101 | *P. a.* | R | R | R | R | S | S | S | S | S | S |
| 38 | *K. p.* | HR | HR | R | R | S | S | S | S | R | R | 102 | *P. a.* | R | R | R | R | S | S | S | S | S | S |
| 39 | *K. p.* | R | R | R | R | R | R | S | S | R | R | 103 | *P. a.* | R | R | R | R | S | S | S | S | S | S |
| 40 | *K. p.* | HR | HR | R | R | HR | HR | S | S | R | R | 104 | *P. a.* | HR | HR | HR | HR | HR | HR | S | S | HR | HR |
| 41 | *K. p.* | HR | HR | R | R | S | S | S | S | R | R | 105 | *P. a.* | HR | HR | HR | HR | S | S | S | S | S | S |
| 42 | *K. p.* | HR | HR | HR | HR | S | S | HR | HR | R | R | 106 | *P. a.* | HR | HR | HR | HR | S | S | S | S | S | S |
| 43 | *K. p.* | R | R | R | R | S | S | S | S | R | R | 107 | *P. a.* | R | R | R | R | S | S | S | S | S | S |
| 45 | *K. p.* | HR | HR | R | R | S | S | S | S | S | S | 108 | *P. a.* | R | R | R | R | S | S | S | S | S | S |
| 46 | *K. p.* | HR | HR | R | R | S | S | S | S | R | R | 109 | *P. a.* | R | R | R | R | S | S | S | S | S | S |
| 47 | *K. p.* | HR | HR | R | R | S | S | S | S | R | R | 110 | *P. a.* | HR | R | R | R | S | S | S | S | S | S |

To further demonstrate their quantitative consistency, we additionally plotted RF_DnD_ versus RF_PAP_ for all isolates against each antibiotic (Supplementary Fig. 3). Across the board, the points followed a linear trend along the line of equality (y=x), demonstrating strong quantitative agreement between the two assays.

### Our assays can quantify persistence

Persistence refers to a phenomenon in which a subpopulation of bacterial cells enters a non-growing or abnormally slow-growing state, allowing them to tolerate otherwise lethal antibiotic treatment ^29^. This tolerant state is transient: once the antibiotic is removed, these persister cells can exhibit normal growth.

Persistence is typically measured using a CFU-based time-kill assay, which exploit this transiency ^29^. In this assay, a bacterial culture was exposed to high concentrations of a bactericidal antibiotic to eliminate actively growing cells. Samples are collected at different time points, washed, and plated on antibiotic-free agar; CFUs reflects the number of surviving persisters. However, like PAP tests, this approach requires manual plating and colony counting, making it labor-intensive and time-consuming.

We therefore evaluated whether our dilution-to-extinction, delay-to-growth, and DnD assays could also quantify persister cells. While these assays were previously performed in the presence of antibiotics to measure resistant cells that grow despite antibiotic exposure, persisters are not resistant and can grow only after the antibiotic is removed. For this reason, traditional persistence assays are conducted in antibiotic-free media following antibiotic exposure. We adopted the same strategy here, performing our assays in antibiotic-free broth after transient antibiotic treatment.

Previous work, including by our group, has shown that exponentially growing *E. coli* cultures contain negligible levels of persisters, whereas overnight stationary-phase cultures exhibit dramatically increased persistence ^7^. Another well-characterized model of enhanced persistence involves deletion of *bfmS* in *Acinetobacter baumannii* ^32^. We applied our assays to both conditions, alongside CFU-based assays for comparison.

Across all cases, our assays accurately quantified persistence frequencies and reproduced the expected biological patterns (Fig. 3a-b). When we determine the deviation of each assay from the CFU-based assay (as done in Fig. 2f), we again found that DnD offers the greatest accuracy and reproducibility.

**Fig. 3:**
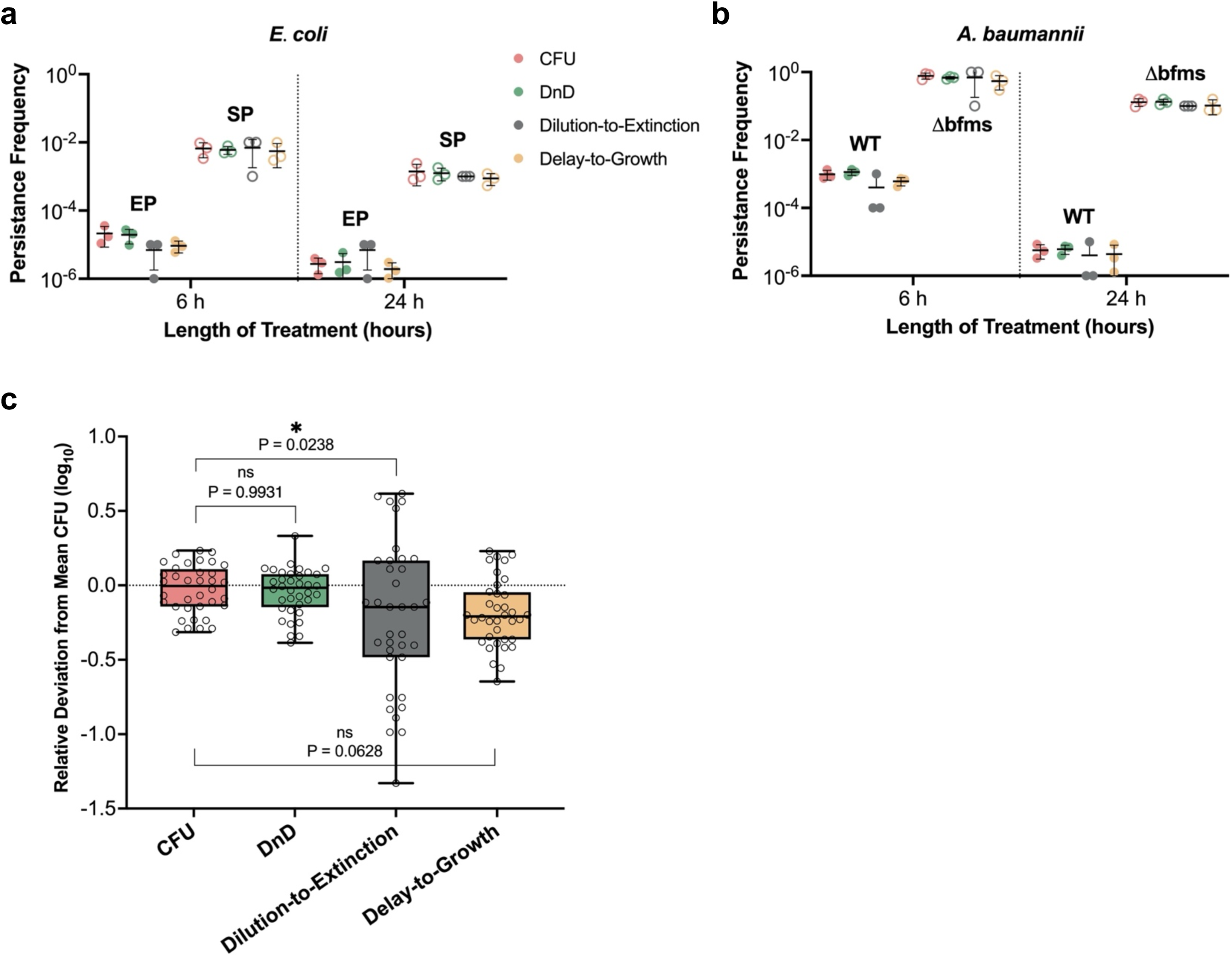
Quantification of bacterial persistence. **a–b.** Persistence frequencies were measured at 6 and 24 hours after antibiotic treatment. Extended incubation in stationary phase in *E. coli* ^7^ and deletion of *bfmS* in *A. baumannii* ^32^ are known to elevate persistence levels. Traditionally, CFU-based assays have been used to quantify persistence. For comparison, we quantified persistence frequencies using dilution-to-extinction, delay-to-growth, and DnD assays. All methods recapitulated the expected biological trends and reproduced persistence frequencies obtained by CFU-based assays. Following published protocols, 100 μg/mL ampicillin for *E. coli* ^7^ and 128 μg/mL carbenicillin for *A. baumannii* ^32,37^ were used. EP: exponential phase; SP: stationary phase **c.** Persistence frequencies determined by each method were statistically compared to CFU-based results, following the same log₁₀ deviation analysis described in Fig. 2d. The DnD assay exhibited the lowest deviation and variance, indicating the highest accuracy and reproducibility among all methods tested.

## Discussion

To address the complex and evolving landscape of antibiotic insensitivity, there is a pressing need for practical, high-resolution susceptibility testing that can be implemented across both research and clinical settings. The DnD assay was developed to meet this need. By leveraging two underutilized experimental axes—cell number and observation time—DnD enables accurate and quantitative detection of rare antibiotic-insensitive cells embedded within otherwise susceptible populations, down to frequencies of 1 in 100 million.

Although PAP provides similarly accurate measurements, it is labor-intensive and time-consuming, requiring 2–3 hours of hands-on time per condition, mostly devoted to agar preparation and manual CFU counting, which is unpractical for routine testing. In contrast, DnD uses a standard broth-based format and requires only 5–15 minutes of setup. Moreover, DnD is readily automatable: all measurements and analysis in this study were performed using a standard plate reader. The most labor-intensive steps—antibiotic media preparation and serial dilution—can be automated using increasingly available robotic liquid handling systems, further reducing the set-up time.

Another strength of the DnD framework is its modularity: its two components—dilution-to-extinction and delay-to-growth—can be used independently, depending on experimental needs or available resources. The dilution-to-extinction assay requires a single visual readout of turbidity, making it suitable for resource-limited settings. Its reliance on 10-fold serial dilutions introduces an intrinsic ∼10-fold uncertainty in RF estimates, as evidenced by a similar degree of deviation in our analysis. This level of precision is sufficient for many applications—particularly when the goal is to determine whether resistant subpopulations are present or not. For greater quantitative resolution, the dilution factor can be reduced (e.g., from 10-fold to 2-fold), though this comes at the cost of exponentially increasing dilution steps.

A more practical strategy for improving precision of the dilution-to-extinction assay is to increase the number of biological replicates. In particular, the most probable number (MPN) analysis method provides a statistical framework for estimating the number of discrete units (e.g., resistant cells) based on the presence or absence of growth in multiple subdivided samples^33^. It was traditionally used to estimate bacterial abundance in environmental samples ^34^. We tested how many replicates are needed to reliably measure RF in the dilution-to-extinction assay. We found that increasing the number of replicates indeed reduces the uncertainty in the RF estimate (Supplementary Fig. 4), although the benefit plateaus beyond five replicates. When we applied the MPN analysis of six replicates to the heteroresistant strains (Fig. 2, blue color), we found that dilution-to-extinction yielded the RF estimates comparable to those from PAP and DnD (Fig. 2d), offering a useful strategy to improve quantitative accuracy in resource-limited settings without specialized equipment.

In contrast, the benefit of the delay-to-growth assay is that it requires only a single culture per condition. Therefore, it is ideal when sample size is limited. It relies on OD monitoring, which can be performed using a basic, widely available spectrophotometer costing less than $1,000. Our data confirm that this method can detect rare resistant subpopulations.

The DnD assay combines these two orthogonal approaches into a unified framework, rigorously and accurately measuring the frequency of antibiotic-insensitive subpopulations—delivering high-resolution susceptibility profiles that standard assays cannot provide. This capability makes DnD a powerful tool for dissecting diverse resistance mechanisms and for detecting the earliest stages of resistance evolution, when intervention is still possible ^35,36^. In clinical settings, such information could guide more informed antibiotic choices and reduce the risk of resistance amplification. For epidemiological surveillance, DnD offers a scalable approach to uncover the true prevalence and diversity of resistance mechanisms across clinical populations. In research, it enables systematic discovery of new resistance phenotypes and mapping their evolutionary trajectories. Together, these strengths position DnD to close a critical gap in antibiotic resistance detection and accelerate both clinical and research advances.

## Data Availability Statements

Source data for figures are provided in the Source Data file.

## Code Availability Statements

No custom-built software was developed.

## Acknowledgements

This work was funded by NIH (1U19AI158080, MM, MK). We thank David Weiss, Jake Choby and Dan Andersson for sharing *Escherichia coli, Enterobacter cloacae* and *Klebsiella pneumoniae* isolates; Sarah Satola for sharing *Pseudomonas aeruginosa* isolates; Phil Rather and Ralph Isberg for sharing *Acinetobacter baumannii* strains.

## Author Contributions

MM and MK conceived the study. MM designed and carried out the experiments. MK secured funding and provided resources. MK wrote the manuscript. All authors read and approved the manuscript.

## Method

### Bacterial strains and culture conditions

Colistin-susceptible and resistant *Enterobacter cloacae* (AMK105 and AMK104 in Fig. 1) were obtained from David Weiss’ lab. Previously characterized heteroresistant strains (AMK107, AMK117, AMK118, AMK120, KMK4 in Fig. 2 and Supplementary Fig. 2) ^17,30,31^ were obtained from David Weiss’ and Dan Andersson’s lab. Clinical isolates (Table 1) were obtained through the Georgia Emerging Infections Program, as part of the CDC’s Multi-site Gram-negative Surveillance Initiative (MuGSI) in Georgia, USA. Wild-type *Acinetobacter baumannii* ATCC7978 and its Δ*bfmS* derivative (KMK111 and KMK112 in Fig. 3) were obtained from Isberg lab at Tufts University ^32^. *Escherichia coli* K-12 strain NCM3722 (referred to as NMK1) was obtained from the Kim Lab bacterial collection ^11^.

Mueller-Hinton broth (MHB; BD Difco #275730) were used. Bacterial cultures were grown in MHB at 37°C with shaking at 250 rpm, in 20 x 150mm borosilicate glass culture tubes (Fisher Scientific #1496133). Exponentially growing cultures, typically at the OD_600_ of 0.1, were used for the assay.

### Antibiotics

Colistin sulfate (#1264-72-8), tobramycin (#32986-56-4), gentamicin sulfate (#1405-41-0) and carbenicillin disodium (#4800-94-6) were obtained from Sigma-Aldrich. Meropenem trihydrate (#119478-56-7) was purchased from Research Product Industry. Tetracycline hydrochloride (#64-75-5) and ampicillin (#69-52-3) were obtained from Bio Basic.

### Population analysis profile (PAP)

PAP was performed as previously described ^30^. Briefly, a single colony of a given strain was inoculated into 2 mL MHB from -80 °C glycerol stock and grown with shaking at 250 rpm. After overnight growth (∼16 hours), cultures were diluted to OD_600_ = 0.001 and grown until OD_600_ reached ∼ 0.1 before being serially diluted in MHB in a 96-well plate (Corning #3598). From each dilution, 5 µL was plated on Mueller-Hinton agar (MHA; BD Difco #225250) containing antibiotics at indicated concentrations. Colonies were enumerated after 24 hours. Resistance frequency (RF) at indicated antibiotic concentration was calculated by dividing the number of colonies on antibiotic-free MHA by the number of surviving colonies on MHA containing antibiotics.

### Dilution-to-Extinction Assay

For resistance frequency (RF) measurements, bacterial cultures were adjusted to approximately 10⁹ CFU/mL based on OD₆₀₀ readings and subjected to 10-fold serial dilutions in Mueller-Hinton Broth (MHB) up to 10⁻⁸. Dilutions were performed across rows in a 96-well microtiter plate (Corning #3596), with each well containing 200 μL of the diluted culture. For each dilution, parallel wells were prepared in MHB with or without indicated concentrations of antibiotics.

Plates were incubated at 37 °C for ∼18 hours. After incubation, wells were visually inspected for turbidity. The first dilution in the series that lacks turbidity defines the extinction point. RF was estimated as the difference in dilution factor between the extinction points in antibiotic-free versus antibiotic-containing conditions.

For persistence frequency measurements, exponential-phase (OD₆₀₀ ∼ 0.1) and 3 days stationary-phase (OD₆₀₀ ∼ 4.0) cultures of *E. coli* NMK1, as well as exponential phase cultures of wild-type and Δ*bfmS A. baumannii*, were exposed to the indicated concentrations of antibiotics for 6 or 24 hours. Following treatment, cultures were washed twice with phosphate-buffered saline (PBS) to remove residual antibiotics and then subjected to 10-fold serial dilutions in 200 μL MHB across rows of 96-well plates, as described above. Plates were incubated at 37 °C for ∼24 hours. After incubation, wells were visually inspected for turbidity. Persistence frequency was calculated by comparing the extinction point of the treated culture to that of the untreated control.

### Delay-to-Growth Assay

For resistance frequency measurements, exponential-phase bacterial cultures (OD₆₀₀ ∼ 0.1) were inoculated into 200 μL Mueller-Hinton Broth (MHB) in a 96-well microtiter plate, with wells containing MHB with and without the indicated concentrations of antibiotics. Plates were incubated at 37 °C with shaking at 250 rpm for 18 to 24 hours. OD₆₀₀ was measured and recorded at 10-minute intervals using a spectrophotometer.

To estimate the number of resistant cells in the original population, the exponential phase of the recovery growth curve in antibiotic-containing wells was fitted. The extrapolated OD₆₀₀ at t = 0 (OD₆₀₀ᴿ) represents the contribution of the resistant subpopulation. This value was normalized to the initial OD₆₀₀ of the total population (OD₆₀₀ᴾ), yielding the resistance frequency (RF): OD₆₀₀ᴿ / OD₆₀₀ᴾ.

For persistence frequency measurements, *E. coli* NMK1 and *A. baumannii* (wild-type and Δ*bfmS*) were treated with the indicated concentrations of antibiotics for 6 or 24 hours. After treatment, cells were washed twice with PBS to remove residual antibiotics, then re-inoculated into 200 μL of fresh MHB in a 96-well microtiter plate. Cultures were incubated at 37 °C with shaking at 250 rpm for ∼24 hours, with OD₆₀₀ monitored at 10-minute intervals.

The OD₆₀₀ of the persister subpopulation (OD₆₀₀^PS^) was estimated by extrapolating the exponential phase of the post-treatment growth curve back to time zero. Persistence frequency was calculated as the ratio of OD₆₀₀^PS^ to the OD₆₀₀ of the total population (OD₆₀₀^P^): PF = OD₆₀₀^PS^/ OD₆₀₀^P^.

### Dilution-and-Delay (DnD) Assay

Exponential-phase (OD_600_ ∼0.1) bacterial cultures were subjected to 10-fold serial dilutions in MHB across rows of a 96-well microtiter plate, with each well containing 200 μL of culture supplemented with or without the indicated concentrations of antibiotics. Plates were incubated at 37 °C with shaking at 250 rpm for approximately 18 hours. OD₆₀₀ was recorded at 10-minute intervals using the microplate reader.

For wells that exhibited detectable growth, the exponential phase of the recovery curve was fitted and extrapolated back to time zero as discussed above to determine RFs. The final RF value was obtained by averaging the RF values across all growing wells in the dilution series.

For persistence frequency measurements, *E. coli* NMK1 and *A. baumannii* (wild-type and Δ*bfmS*) were exposed to the indicated concentrations of antibiotics for 6 or 24 hours at 37 °C with shaking. After treatment, cultures were washed twice with fresh MHB to remove residual antibiotics, then serially diluted in 200 μL fresh MHB across rows of a 96-well plate as described above. Plates were incubated at 37 °C with shaking at 250 rpm, and OD₆₀₀ was recorded every 10 minutes for ∼24 hours.

### CFU-based Time-kill Assay

Exponential-phase (OD₆₀₀ ∼ 0.1) and 3 days stationary-phase (OD₆₀₀ ∼ 4) cultures of *E. coli* NMK1, and *A. baumannii* (wild-type and Δ*bfmS*) were exposed to the indicated concentrations of antibiotics for 6 or 24 hours at 37 °C with shaking. After treatment, 100 μL aliquots were removed from the cultures, washed twice with PBS to remove residual antibiotics, serially diluted in PBS, and plated on MHA. Plates were incubated at 37 °C for ∼24 hours, and colony-forming units (CFU) were enumerated to determine viable cell counts. To calculate persistence frequency, the number of surviving cells at 6 or 24 hours was divided by the initial CFU count (t = 0) prior to antibiotic exposure.

### Statistical analysis

Statistical analysis was conducted using GraphPad Prism. Details of biological replicates and statistical tests are provided in the corresponding figure legends. All data sets were tested for normality using the Shapiro-Wilk test and were confirmed to meet the normality criteria. Statistical analyses were performed as appropriate based on the experimental design.

## References

1 Martens, E. & Demain, A. L. The antibiotic resistance crisis, with a focus on the United States. The Journal of Antibiotics 70, 520 (2017). 10.1038/ja.2017.30

2 Currie, C. J. et al. Antibiotic treatment failure in four common infections in UK primary care 1991-2012: longitudinal analysis. BMJ 349 (2014). 10.1136/bmj.g5493

3 Blair, J. M. A., Webber, M. A., Baylay, A. J., Ogbolu, D. O. & Piddock, L. J. V. Molecular mechanisms of antibiotic resistance. Nat Rev Micro 13, 42–51 (2015). 10.1038/nrmicro3380

4 Band, V. I. et al. Colistin Heteroresistance Is Largely Undetected among Carbapenem-Resistant Enterobacterales in the United States. mBio 12, e02881–02820 (2021). doi:10.1128/mBio.02881-20

5 Nicoloff, H., Hjort, K., Levin, B. R. & Andersson, D. I. The high prevalence of antibiotic heteroresistance in pathogenic bacteria is mainly caused by gene amplification. Nature Microbiology 4, 504–514 (2019). 10.1038/s41564-018-0342-0

6 Balaban, N. Q., Merrin, J., Chait, R., Kowalik, L. & Leibler, S. Bacterial persistence as a phenotypic switch. Science 305, 1622–1625 (2004). 10.1126/science.1099390

7 Şimşek, E. & Kim, M. Power-law tail in lag time distribution underlies bacterial persistence. Proceedings of the National Academy of Sciences 116, 17635–17640 (2019). 10.1073/pnas.1903836116

8 Sánchez-Romero, M. A. & Casadesús, J. Contribution of phenotypic heterogeneity to adaptive antibiotic resistance. Proceedings of the National Academy of Sciences 111, 355–360 (2014). 10.1073/pnas.1316084111

9 Motta, S. S., Cluzel, P. & Aldana, M. Adaptive Resistance in Bacteria Requires Epigenetic Inheritance, Genetic Noise, and Cost of Efflux Pumps. PloS one 10, e0118464 (2015). 10.1371/journal.pone.0118464

10 Coates, J. et al. Antibiotic-induced population fluctuations and stochastic clearance of bacteria. eLife 7, e32976 (2018). 10.7554/eLife.32976

11 Stine, W., Akiyama, T., Weiss, D. & Kim, M. Lineage-dependent variations in single-cell antibiotic susceptibility reveal the selective inheritance of phenotypic resistance in bacteria. Nature communications 16, 4655 (2025). 10.1038/s41467-025-59807-x

12 Dewachter, L., Fauvart, M. & Michiels, J. Bacterial Heterogeneity and Antibiotic Survival: Understanding and Combatting Persistence and Heteroresistance. Mol. Cell 76, 255–267 (2019). 10.1016/j.molcel.2019.09.028

13 Akiyama, T. & Kim, M. Stochastic response of bacterial cells to antibiotics: its mechanisms and implications for population and evolutionary dynamics. Current opinion in microbiology 63, 104–108 (2021). 10.1016/j.mib.2021.07.002

14 Band, V. I. & Weiss, D. S. Heteroresistance to beta-lactam antibiotics may often be a stage in the progression to antibiotic resistance. PLOS Biology 19, e3001346 (2021). 10.1371/journal.pbio.3001346

15 Levin-Reisman, I. et al. Antibiotic tolerance facilitates the evolution of resistance. Science 355, 826–830 (2017). doi:10.1126/science.aaj2191

16 Windels, E. M. et al. Bacterial persistence promotes the evolution of antibiotic resistance by increasing survival and mutation rates. The ISME journal 13, 1239–1251 (2019). 10.1038/s41396-019-0344-9

17 Pal, A. & Andersson, D. I. Bacteria can compensate the fitness costs of amplified resistance genes via a bypass mechanism. Nature communications 15, 2333 (2024). 10.1038/s41467-024-46571-7

18 CLSI. (ed Clinical and Laboratory Standards Institute) (2025).

19 (EUCAST), E. C. f. A. S. T. Determination of minimum inhibitory concentrations (MICs) of antibacterial agents by broth dilution. Clinical Microbiology and Infection 9, ix-xv (2003). 10.1046/j.1469-0691.2003.00790.x

20 Kowalska-Krochmal, B. & Dudek-Wicher, R. The Minimum Inhibitory Concentration of Antibiotics: Methods, Interpretation, Clinical Relevance. Pathogens 10 (2021). 10.3390/pathogens10020165

21 Matuschek, E., Brown, D. F. J. & Kahlmeter, G. Development of the EUCAST disk diffusion antimicrobial susceptibility testing method and its implementation in routine microbiology laboratories. Clinical Microbiology and Infection 20, O255–O266 (2014). 10.1111/1469-0691.12373

22 Turnidge, J. & Paterson, D. L. Setting and revising antibacterial susceptibility breakpoints. Clin Microbiol Rev 20, 391–408, table of contents (2007). 10.1128/cmr.00047-06

23 El-Halfawy, O. M. & Valvano, M. A. Antimicrobial Heteroresistance: an Emerging Field in Need of Clarity. Clinical Microbiology Reviews 28, 191–207 (2015). 10.1128/cmr.00058-14

24 Band, V. I. & Weiss, D. S. Heteroresistance: A cause of unexplained antibiotic treatment failure? PLoS pathogens 15, e1007726 (2019). 10.1371/journal.ppat.1007726

25 Andersson, D. I., Nicoloff, H. & Hjort, K. Mechanisms and clinical relevance of bacterial heteroresistance. Nature Reviews Microbiology 17, 479–496 (2019). 10.1038/s41579-019-0218-1

26 Baltekin, Ö., Boucharin, A., Tano, E., Andersson, D. I. & Elf, J. Antibiotic susceptibility testing in less than 30 min using direct single-cell imaging. Proceedings of the National Academy of Sciences 114, 9170–9175 (2017). doi:10.1073/pnas.1708558114

27 Deris, B. et al. The innate growth bistability of antibiotic resistant bacteria. Science 342, 1237435–1237431 (2013, * equal contribution).

28 Ronneau, S., Hill, P. W. S. & Helaine, S. Antibiotic persistence and tolerance: not just one and the same. Current opinion in microbiology 64, 76–81 (2021). 10.1016/j.mib.2021.09.017

29 Balaban, N. Q. et al. Definitions and guidelines for research on antibiotic persistence. Nature Reviews Microbiology 17, 441–448 (2019). 10.1038/s41579-019-0196-3

30 Band, V. I. et al. Antibiotic failure mediated by a resistant subpopulation in Enterobacter cloacae. Nature Microbiology 1, 16053 (2016). 10.1038/nmicrobiol.2016.53

31 Pereira, C., Larsson, J., Hjort, K., Elf, J. & Andersson, D. I. The highly dynamic nature of bacterial heteroresistance impairs its clinical detection. Communications Biology 4, 521 (2021). 10.1038/s42003-021-02052-x

32 Geisinger, E., Mortman, N. J., Vargas-Cuebas, G., Tai, A. K. & Isberg, R. R. A global regulatory system links virulence and antibiotic resistance to envelope homeostasis in Acinetobacter baumannii. PLoS pathogens 14, e1007030 (2018). 10.1371/journal.ppat.1007030

33 Chandrapati, S. & Williams, M. G. in Encyclopedia of Food Microbiology (Second Edition) (eds Carl A. Batt & Mary Lou Tortorello) 621-624 (Academic Press, 2014).

34 Rowe, R., Todd, R. & Waide, J. Microtechnique for Most-Probable-Number Analysis. Applied and environmental microbiology 33, 675–680 (1977). doi:10.1128/aem.33.3.675-680.1977

35 Band, V. I. et al. Antibiotic combinations that exploit heteroresistance to multiple drugs effectively control infection. Nat Microbiol 4, 1627–1635 (2019). 10.1038/s41564-019-0480-z

36 Baym, M., Stone, L. K. & Kishony, R. Multidrug evolutionary strategies to reverse antibiotic resistance. Science 351, aad3292 (2016). 10.1126/science.aad3292

37 Bhargava, N., Sharma, P. & Capalash, N. Pyocyanin stimulates quorum sensing-mediated tolerance to oxidative stress and increases persister cell populations in Acinetobacter baumannii. Infect Immun 82, 3417–3425 (2014). 10.1128/iai.01600-14

