## Supplementary Material for "High-Resolution Detection of Hidden Antibiotic Resistance with the Dilution-and-Delay (DnD) Susceptibility Assay"

### Supplementary Figures

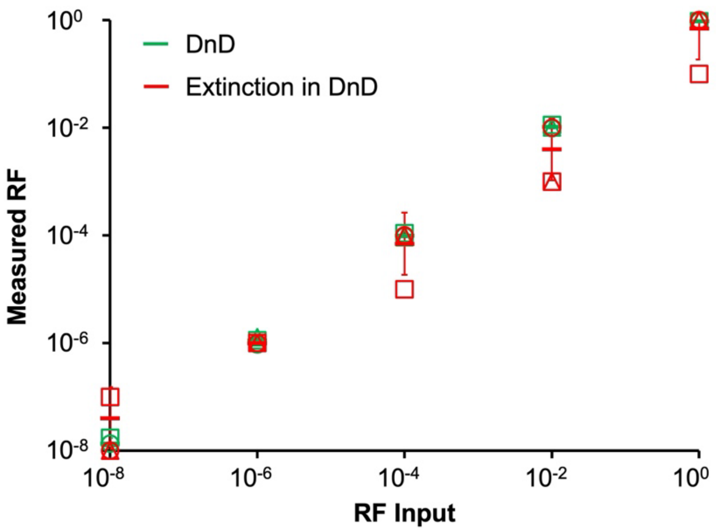


**Supplementary Fig. 1. Validation of the Dilution-and-Delay (DnD) assay using defined input mixtures.**

As described in the main text, a colistin-susceptible strain (AMK105) was mixed with a fully resistant strain (AMK104) at defined input frequencies. Resistance frequencies (RFs) were measured. Green symbols indicate RFs measured from the wells showing growth recovery (adopted from Fig. 1f). Red symbols indicate RFs from extinction points. Their RFs were within a 10-fold range, confirming the internal consistency. Each experiment was performed in biological triplicate. Individual replicates are shown as different symbols, with lines and error bars indicating the mean and standard deviation.

**a**

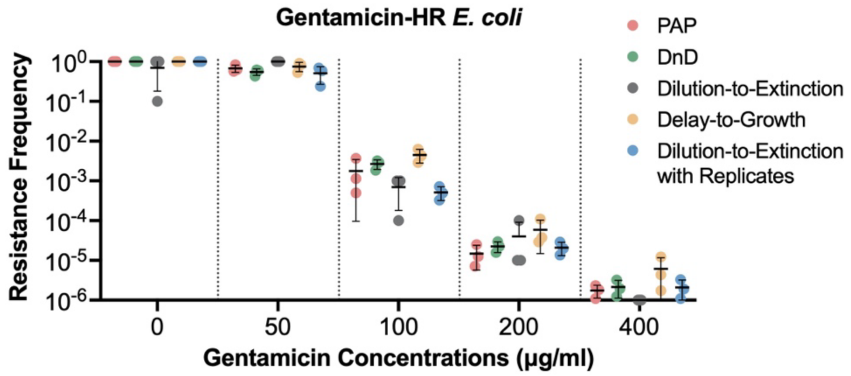


**b**

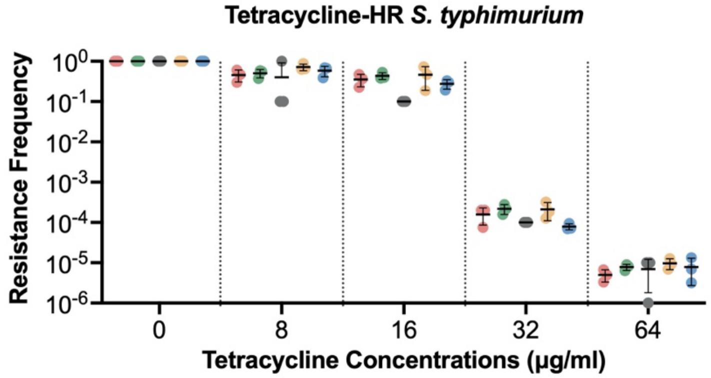


**Supplementary Fig. 2: Validation of the DnD assay against heteroresistant clinical isolates.**

**a–b.** In addition to the three heteroresistant (HR) strains shown in Fig. 2, resistance frequencies (RF) were measured for two additional clinical HR isolates: gentamicin-HR *E. coli* (AMK118) and tetracycline-HR *S. typhimurium* (AMK120). Each experiment was performed in biological triplicate. Individual replicates are shown as different symbols, with black lines and error bars indicating the mean and standard deviation. All three assays recapitulated PAP-derived RF, $RF_{pap}$, across antibiotics and concentrations.

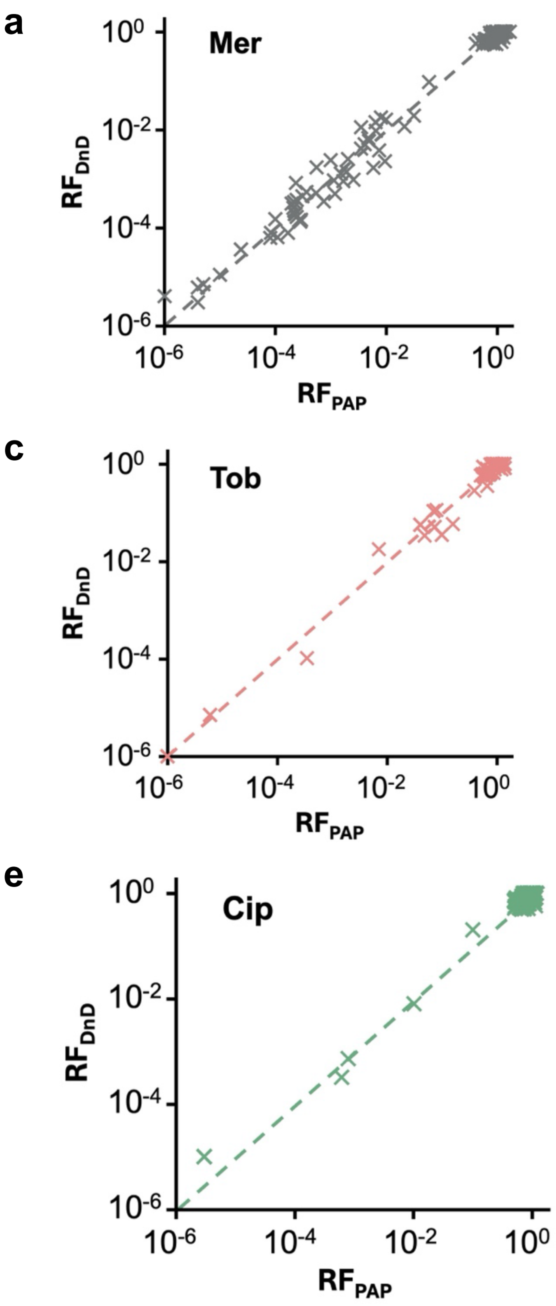


**Supplementary Fig. 3: Comparing resistance frequency RF measured using the DnD ($RF_{DnD}$) and PAP ($RF_{PAP}$) across clinical isolates.**

**a-e.** $RF_{DnD}$ measured in Table 1 were plotted against $RF_{PAP}$ for each antibiotic: a) meropenem (Mer), b) piperacillin/tazobactam (Pip/Tazo), c) tobramycin (Tob), d) colistin (Col), and **e)** ciprofloxacin (Cip). The diagonal dashed line denotes y = x (the line of equality).

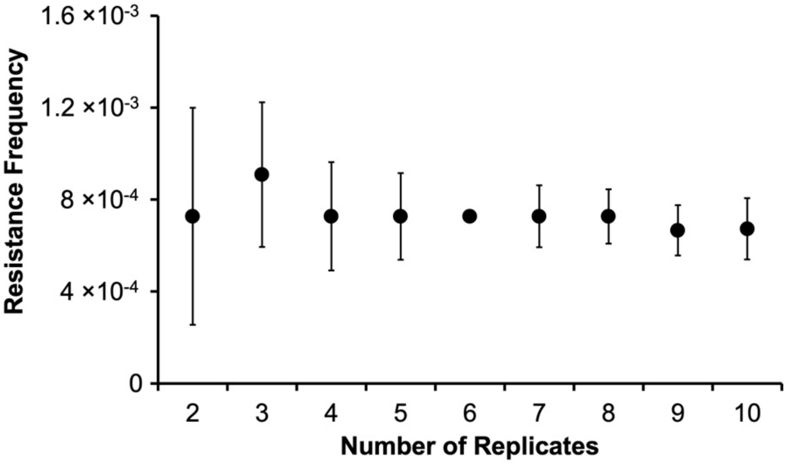


**Supplementary Fig. 4. Increasing the number of replicates improves the precision of dilution-to-extinction assays.**

Resistant frequencies (RF) were calculated using most probable number (MPN) analysis applied to dilution-to-extinction data with varying numbers of biological replicates. Increasing the number of replicates reduced the uncertainty (as reflected by narrower confidence intervals), indicating the use of MPN-based analysis with a moderate number of replicates (e.g., $n \geq 5$) as a practical approach to improve RF quantification.

**Supplementary Table 1. Minimum inhibitory concentrations (MICs) of the five previously characterized heteroresistant bacterial strains tested in Fig. 2 and Supplementary Fig. 2.**

| Strain | Species | Antibiotic Tested | MIC (μg/mL) |
|---|---|---|---|
| AMK107 | *E. cloacae* | Colistin | 512 |
| AMK117 | *E. coli* | Tobramycin | 40 |
| AMK118 | *E.coli* | Gentamicin | 400 |
| AMK120 | *S. typhimurium* | Tetracycline | 64 |
| KMK4 | *K. pneumoniae* | Meropenem | 32 |
